# Molecular architecture of Influenza A virions

**DOI:** 10.64898/2026.04.02.715802

**Authors:** Swetha Vijayakrishnan, Jack C. Hirst, Sarah Cole, Svenja S. Hester, Vattipally B. Sreenu, Colin Loney, Wael Kamel, Roman Fischer, Terry K. Smith, Ludovic Autin, David Bhella, Edward Hutchinson

## Abstract

Influenza A viruses (IAV) are clinically important pathogens that cause seasonal epidemics and pandemics in humans. IAV produce pleomorphic, enveloped virions, which can range from a spherical or bacilliform morphology, the predominant form in the most commonly studied laboratory strains, to long filamentous virions which are characteristic of clinical and veterinary isolates. Understanding the structure and function of filamentous virions is crucial for clarifying their role in viral persistence and immune evasion, and for informing the development of therapeutics that target their entry and/or egress pathways.

Structural characterisation of influenza virions is challenging however owing to their fragility, heterogeneity and compared to most virus particles, unusually large size. Here, we combined structural and compositional approaches with integrative modelling to define the complete molecular architecture of influenza virions. In doing so we provide the first description of distinctive structural features of IAV filaments, including the selective incorporation of lipids, specific enrichment of viral and host proteins, and a viral cytoskeleton including a secondary helical layer within the viral capsid and extended fibrils of cofilactin. Collectively our findings suggest an important regulatory role for cofilactin in driving filament morphogenesis and provide important insights into the organisation and composition of IAV filamentous virions.

## Introduction

Influenza A viruses (IAV) are among the most important respiratory pathogens affecting humans worldwide. Each year they cause seasonal epidemics, infecting millions of people, and occasionally triggering pandemics of far greater impact that occur after the emergence of new reassorting viruses. Globally IAV infect around 10% of the population annually, causing about half a million deaths, with the heaviest burden falling on low- and middle-income countries^1,2^. Although IAV infect people of all ages, young children, pregnant women, and the elderly (> 65 years) are most at risk of severe illness and death^3,4^.

IAV produce enveloped virions that contain a negative-sense, segmented RNA genome. The viral envelope is embedded with three proteins critical for infection: haemagglutinin (HA), which mediates receptor binding and entry; neuraminidase (NA), which facilitates viral release; and the ion channel M2, which supports uncoating^5-9^. Beneath the lipid envelope lies the matrix protein M1 that forms a protein endoskeleton, providing structural support and regulating assembly^10^. The interior of the virion contains the genome where the RNA molecules are encapsidated by multiple copies of nucleoprotein (NP) to form right-handed, supercoiled ribonucleoprotein complexes (RNPs)^11^. The RNPs are packed as eight segments arranged in an orderly fashion with a central segment surrounded by seven others in a “7+1” manner, usually lying along the long axis of the virion ^12-14^. High-resolution structural studies using X-ray crystallography, cryo-electron microscopy (cryo-EM), and cryo-electron tomography (cryo-ET) have provided atomic models of trimeric HA^15,16^, tetrameric NA^17,18^, helical M1^19,20^ and RNPs^11,21-23^, deepening our understanding of IAV biology at the protein level. While the structures of individual proteins are either known or can be predicted, their arrangement within the IAV virion remains difficult to define.

Much of our understanding of IAV biology has been derived from studies of virus infection in cell culture, particularly well characterised laboratory adapted strains that in some cases were originally isolated and passaged in animal models. IAV are highly pleomorphic, exhibiting various strain-dependent morphologies^24^. Lab-adapted stains produce spherical and bacilliform virions, typically between 120 and 250 nm in length^25,26^. In contrast, clinical isolates are characterised by the production of strikingly long filamentous virions, often several microns in length^14,27,28^. However structural and mechanistic characterisation of filamentous virions has proven challenging as they are fragile and easily destroyed by standard laboratory purification methods. To overcome this limitation, we previously developed methods for cryo-ET imaging of IAV filamentous virions that preserved their native ultrastructure by propagating the virus in cells grown directly on electron microscopy grids^14^. This approach allowed detailed visualisation of these fragile virions in near-native conditions and enabled us and others to develop a structural understanding of the filamentous phenotype^14,19,29,30^.

M1 and M2 have shown to strongly influence virion morphology, for example a single mutation K102A in M1 of influenza A/WSN/33 renders this normally spherical virion filamentous, or a set of five combined mutations in the C-terminal amphipathic helix of M2 impairs production of filamentous Udorn virus^31-35^. Interestingly, filament formation is also highly dependent on cell type, with polarised epithelial cells producing greater numbers of IAV filaments than non-polarised cells when infected with the same virus strain^33,36-38^. Proteomics studies have shown that IAV incorporates numerous host proteins into budding virions^39,40^, including actin. Although actin is not required for the release of spherical particles, integrity of the actin cytoskeleton is important for filamentous virion budding^41^. Disruption of this network alters lipid-raft domains on the plasma membrane where IAV assembly occurs, correlating with a marked reduction in filamentous virion production^42^. Moreover, actin has been shown to associate with M1 and RNPs, and to modulate HA dynamics, suggesting a coordinated interplay of IAV proteins and the actin cytoskeleton^43,44^. Actin-associated proteins such as myosin and cofilin have also been shown to play critical roles in IAV assembly^45-47^.

Understanding the morphology, organisation and structure of IAV filaments is important to elucidate their functional role during infection and help inform development of effective therapeutics. Although structures of several individual IAV proteins are known, placing them in the context of the complete filamentous virion continues to be challenging owing to the virus’s pleomorphism. To address this, we employed an integrative, multidisciplinary approach combining mass spectrometry, cryo-ET, and computational integrative modelling to generate a comprehensive molecular model of the IAV filaments.

Using comparative multi-omics analysis of spherical versus filamentous virions we revealed distinct compositional differences in their lipid and protein profiles. Filaments contain reduced levels of curvature-promoting lipids, which may contribute to their elongated shape, and show an altered protein composition, with less NA compared to spherical particles. Using cryo-ET imaging of IAV filaments budding directly from cells, we examined fibrillar densities running along their axes, which were not detected in spherical virions. Using sub-tomogram averaging, we reconstructed these fibrils at 12 Å resolution, revealing features consistent with them being cofilactin, actin filaments decorated with cofilin. By docking AlphaFold-predicted structures of canine actin and cofilin into our density maps, we built a model of cofilactin that provides a view of its orientation within the filamentous virion and when assembled into higher-order helical bundle assemblies. We then used confocal microscopy and Western blotting to show that IAV infection upregulates cofilin expression in infected cells while promoting its dephosphorylation. Together, these data suggest that IAV may regulate cofilin activity during virion assembly and budding.

Our cryo-ET data also identified a second helical density within filamentous virions that is concentric to the matrix protein (M1) layer. Comparisons to other studies suggest that this is likely to be a secondary M1 helix.

By integrating structural and compositional data we present the first comprehensive integrative molecular model of filamentous IAV, underlining the unique, complex, multilayered organisation of the filamentous virion. Together, our findings highlight the pleomorphic and distinctive molecular organisation of influenza virions, underscoring the importance of host–virus interactions in shaping virion architecture and opening new avenues for targeted therapeutics.

## Results

### Purification of IAV produced limited number of filamentous particles

To obtain a clear understanding of the structure and composition of IAV filamentous virions we carried out large-scale preparations of purified influenza virions. We had previously demonstrated that long IAV filaments (>3 μm) are highly susceptible to damage during purification^14^. Light microscopy suggested a reduced number of long filaments (>4 μm) when iodixanol gradients were used instead of sucrose, although their structural integrity could not be confirmed due to the method’s low resolution^48^. To assess this more precisely, we reanalysed our previously published data of purified A/Udorn/72 IAV (A/Udorn/72) from MDCK cells that was examined by cryo-ET (Fig. 1A)^14^. These samples contained particles of varying morphologies, including bacilliform virions (∼95 nm diameter), short filaments (1–1.5 μm long, 75–80 nm diameter) with occasional varicosities, but rarely filaments exceeding 2 μm. Closer inspection revealed distinct morphological differences between capsule-shaped bacilliform particles and narrower filaments, consistent with prior observations^14^. Longitudinal magnified views frequently showed three RNP segments (yellow arrowheads) lying side-on in both bacilliform (blue circle, Fig. 1A) and short filamentous (black rectangle, Fig. 1A) particles. Quantitative analysis on 127 virus particles using frequency distribution revealed that purified preparations predominantly consisted of bacilliform particles (150–200 nm long), whereas filaments (600 nm–2 μm long) represented only 25.9% of the population (Fig. 1B). While some filaments displayed sparse internal clumps of density, many exhibited fibrillar structures running along their length (white rectangle, Fig. 1A), morphologically distinct from RNPs. However, these tomograms of purified IAV were collected using a 200 kV TEM fitted with a CCD detector and lacked sufficient resolution to resolve and interpret these fibrils clearly.

**Figure 1.**
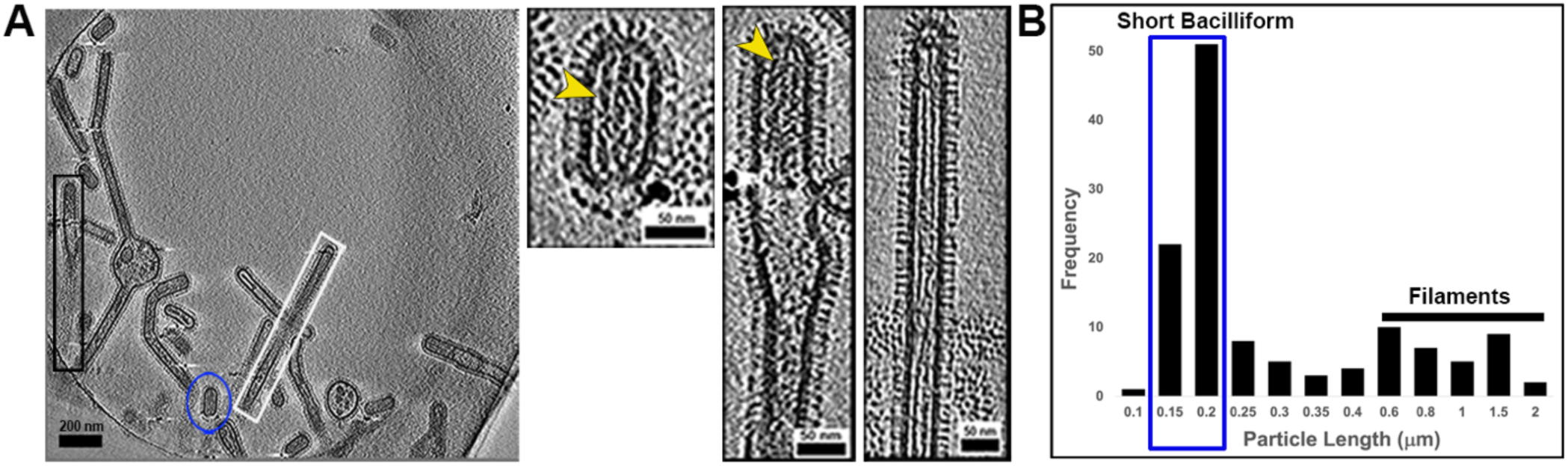
IAV filament distribution in purified virus preparations. Previously published data^14^ of purified A/Udorn/72 from MDCK cells were reanalysed and showed a marked reduction in filaments longer than 2 μm. **(A)** Cryo-ET revealed morphological differences between bacilliform particles and short filaments. Magnified views of bacilliform (blue circle) and filamentous (black rectangle) particles revealed side on arrangement of three RNPs (yellow arrowheads) within virions. Some filaments appear sparsely populated, while others contained distinct internal fibrillar densities (white rectangle) extending along their length. **(B)** Particle length-based frequency distribution analysis demonstrated that bacilliform particles (150-200 nm, blue rectangle) were far more abundant than filamentous particles (0.6-2 μm length) in purified preparations of IAV.

### IAV filaments show distinct composition to spherical virions

It had previously been challenging to analyse the composition of filamentous influenza virions due to their heterogeneous and fragile structures^33,48,49^. To address this, we previously determined that ultracentrifugation through a carefully designed, iso-osmotic density gradient can separate influenza virions by length (Figs. 2A, 2B)^50^. The lengths of a population of influenza virions are typically heterogeneous and vary continuously between short bacilliform virions and elongated filaments (Fig. 1). Due to this, density gradient centrifugation does not clearly separate virions into bacilliform (and spherical) and filamentous populations, but it does concentrate the bacilliform virions in the denser parts of the gradient. As a result, gradient fractions of clearly separated densities allowed us to compare the composition of virion populations that were majority bacilliform or majority filamentous, with the majority of NP protein (which encapsidates the viral genome), and infectivity, found in the bacilliform-enriched fractions (Figs. 2C, 2D).

**Figure 2.**
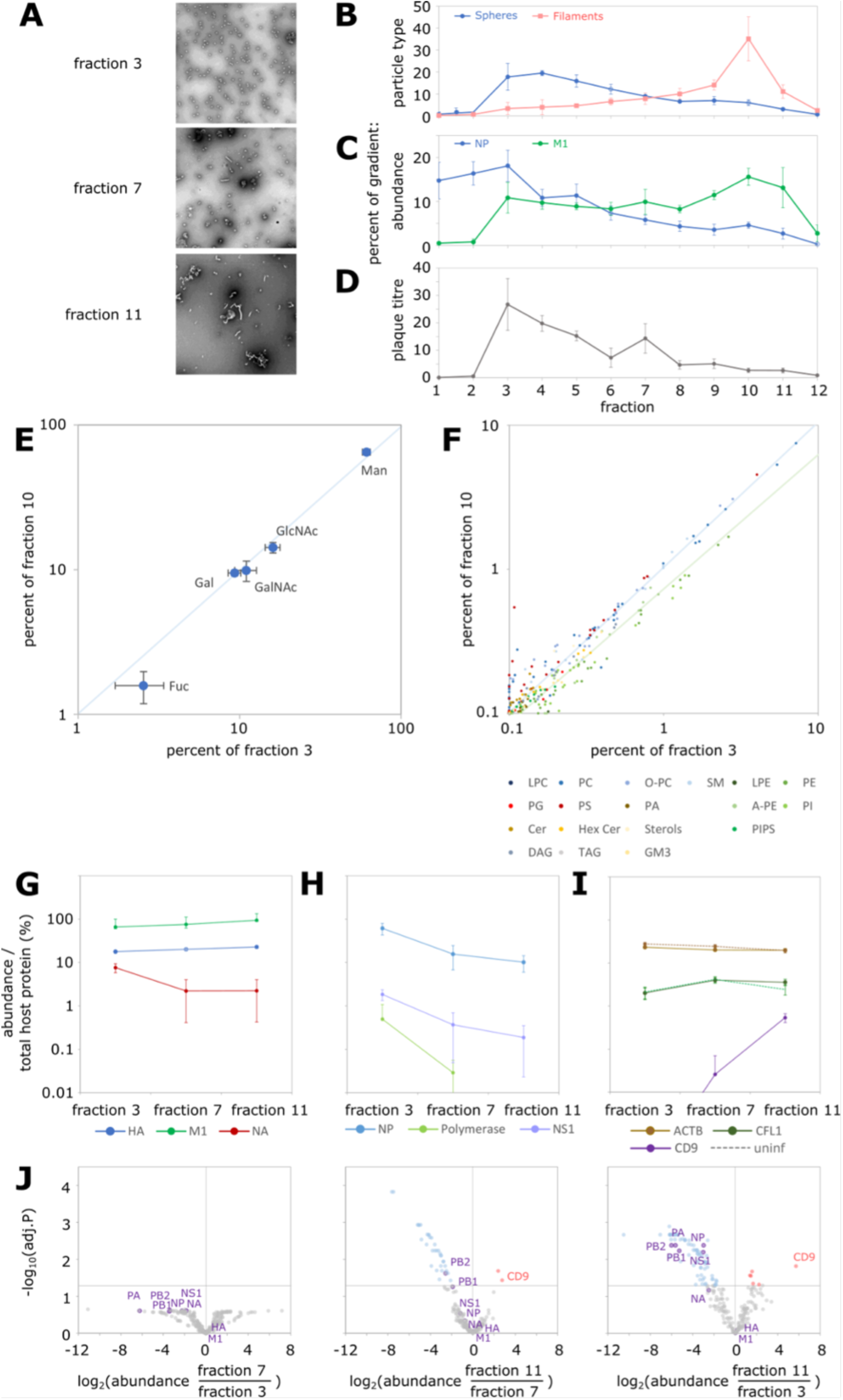
Composition of Influenza Virions of Different Morphologies. Influenza A//Udorn/72 virus was grown in MDCK cells and virions were pelleted through a 10% cushion of iodixanol and then separated on a 20-30% iodixanol gradient overnight by ultracentrifugation at 4 °C and 210,000 *g*. The gradient was separated into 12 fractions, harvested from the bottom of the tube. **(A)** Representative negative stain electron micrographs of virions from fractions 3 (mainly spherical/bacilliform), 7 (intermediate) and 11 (mainly filamentous). **(B)** Virions from each fraction were adsorbed onto glass and virion morphology determined by immunofluorescence. Virions were scored as spherical (including bacilliform) or filamentous as described previously^48^. **(C)** Samples of each fraction were analysed by Western blot for the viral proteins NP and M1. **(D)** The infectious titre of each fraction was determined by plaque assay on MDCK cells. **(E)** Carbohydrates were extracted from the indicated fractions and their abundance analysed by GC-MS. **(F)** Lipids were extracted from the indicated fractions and their abundance analysed by mass spectrometry. **(G)** – **(J**) Proteins were extracted from the indicated fractions, subject to tryptic digest and analysed by LC-MS/MS. The abundance of selected proteins is shown in **(G) – (I)** and the fold changes of all proteins are shown in **(J)**. For **(B)** – **(J)** data from three independent experiments are shown (except for uninfected data in **(I**), for which data are from a single experiment; CD9 was not detected in samples from uninfected cells), with the means and standard deviations shown for **(B)** – **(I**). For **(G) – (I)** an arbitrary value (10^-6^ units) has been added to all data toallow use of zero data on a logarithmic scale.

Using mass spectrometry, we first analysed the glycome of ‘bacilliform’ and ‘filamentous’ fractions (fractions 3 and 10, respectively) and found that the ratios of monosaccharide content were consistent in both fractions (Fig. 2E). We next used electrospray ionisation mass spectrometry to analyse the lipidome of the two fractions (Fig. 2F). We detected over 200 lipid species, the majority of which maintained a consistent ratio between the two fractions. The exceptions were phosphatidylethanolamine (PE) and phosphatidylinositol (PI) class lipids, which were consistently depleted in filamentous virions to 60 % of the levels found in bacilliform virions (Fig. 2F). These lipid classes support membrane fluidity and curvature^51,52^, and it is notable that for filamentous virions most of the membrane surface only curves around one axis, as opposed to curvature around two axes for smaller virions.

Finally, we analysed the virion proteome by mass spectrometry (Figs. 2G – 2J). Considering viral proteins, we observed that HA and M1 are present at similar abundance in fractions across the gradient (Fig. 2G). This was expected, as these proteins densely occupy the virion membrane across its entire surface and so should maintain their ratio regardless of virion morphology.

We also observed a specific depletion of NA in filamentous virions (Figs. 2G, 2J), consistent with structural biology reports indicating that clusters of NA are less densely incorporated into filamentous virions than into bacilliform virions, potentially clustering near one tip^12,24,53^. The proteins that encapsidate the viral genome (PB2, PB1, PA and NP) maintained a consistent ratio to each other and were depleted in filamentous virions to an even greater extent than NA (Figs. 2H, 2J). This is consistent with structural biology reports that filaments, despite their greater length, incorporate at most one copy of the viral genome and often none at all^14,24,53^.

Consistent with our previous studies, we detected NS1 in virions^39^. We had assumed that NS1 might be passively incorporated into virions given its high abundance in the cytoplasm of infected cells at the time of virion budding^54^ – if this were the case it would maintain a consistent ratio with HA and M1. Instead, NS1 was depleted in filamentous virions, maintaining a consistent ratio with the genome-encapsidating proteins (Figs. 2H, 2J). The consistent ratio of NS1 to these proteins suggests that there may be a specific association of NS1 with the viral genome.

Corroborating our previous studies, we detected many host proteins in the virions^39^. Most of these had a similar abundance across all fractions and were also present at similar levels in material from uninfected cells, making their association with virions unclear. An interesting exception was the tetraspanin CD9, which was enriched in filamentous fractions and (despite being common in microvesicles) was undetectable in fractions prepared from the media of uninfected cells (Figs. 2H, 2J). We previously found that CD9 is enriched in the membranes of virions when influenza viruses are grown in mammalian cell culture, but not when they are grown in embryonated chicken eggs^39^, and it is notable that eggs are a culture system which is thought to be unfavourable to the growth of filamentous virions^26^.

Although most host proteins appeared to have similar concentrations in virions of different morphologies, their ability to form complexes with each other would be limited by the dimensions of the virion. To examine this, we required structural information about the virus particles.

### Cryo-ET reveals unique helical structural features within budding IAV filaments

Having established the molecular composition of IAV filaments, we next examined their structural organisation at higher resolution using cryo-ET. As described previously, tilt-series were collected from frozen-hydrated preparations of IAV-infected MDCK cells, propagated directly on quantifoil (R2/2) gold EM grids to preserve the fragile morphology of long (>2 μm) filaments^14^. This approach allowed imaging of virions budding directly from cell edges and 42 tilt-series of undamaged, straight filament segments were acquired. Low-magnification grid views guided the selection of regions of interest (ROIs) where filaments extended from cells (supplementary Fig. S1). These showed well-preserved long filaments exceeding 3 μm in length (Figs. 3A–3C). Selected straight segments (pink boxes) of long filaments in Figs 3A-3C were used for tilt-series acquisition, and tomographic reconstructions revealed ordered virion structures with glycoprotein spikes, a lipid envelope, and additional densities distinct from RNPs (Figs. 3D–3F; supplemental movie S1). Examples of such filament-specific features are shown in Fig. 3G-3K. Among the analysed filamentous virions, 44.5% lacked internal density, 4.8% contained sparse indistinct material (Fig. 3G), 15.6% enclosed extensive internal fibrillar density (Figs. 3D, 3F, 3H; supplemental movie S1), and 36.1% displayed a second inner helix lining the interior surface of the M1 layer but having a longer pitch (Figs. 3E, 3F, 3I, 3J, 3K; supplemental movie S1). Both latter features were clearly distinct from density associated with RNPs (Fig. 3L).

**Figure 3.**
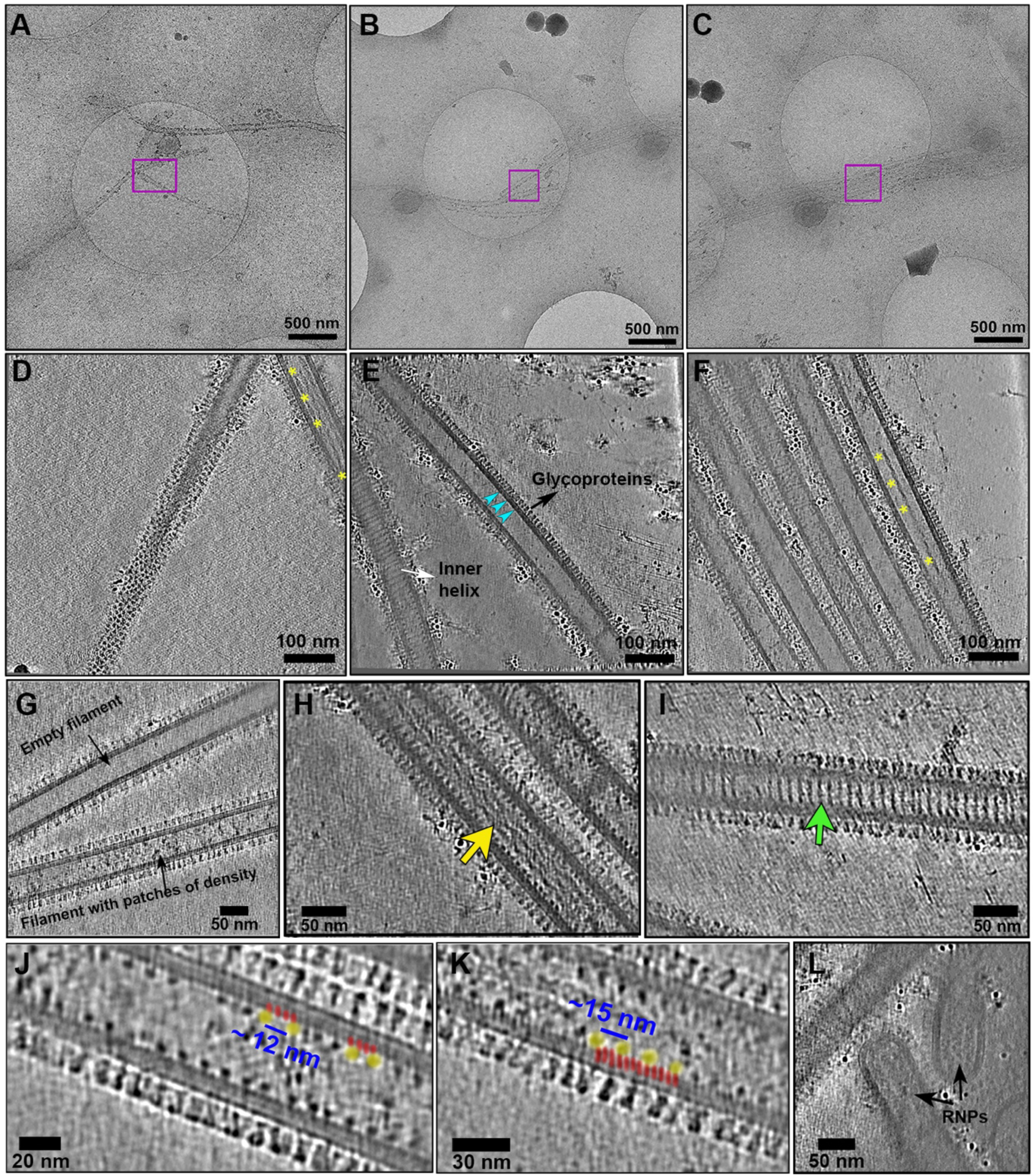
Cryo-ET reveals unique structural features in IAV filaments. **(A, B, C)** Low-magnification imaging of budding filamentous virions identified intact straight segments (pink squares) for tilt-series acquisition. **(D, E, F)** Denoised tomographic slices of these regions showed well-preserved filament architecture with glycoprotein spikes (black arrows) decorating the envelope and supported by the helical M1 scaffold (cyan arrowheads). In addition to these canonical structures, filaments often contained distinct internal features, including **(D, F)** fibrillar densities (yellow stars) and **(E, F)** a secondary inner helix (white arrow). Cryo-ET revealed substantial structural heterogeneity among budding IAV filaments. **(G)** While many appeared empty, others contained patchy internal densities, **(H)** fibrillar structures (yellow arrow), **(I)** a secondary inner helix (green arrow), or both. The inner helix (yellow circles) showed a larger spacing than M1 (red lines), **(J)** corresponding to ∼12 nm (three M1 turns) or **(K)** ∼15 nm (four M1 turns) per turn. Highlighted yellow circles and red lines are shown only for a small section of the filament for ease of visualisation. **(L)** Filaments containing RNPs (black arrows) were rarely detected.

### IAV filaments exhibit a complex ordered envelope

To gain deeper insight into the architecture of IAV filaments, we analysed our tomograms, which revealed that filamentous particles possess a highly ordered and multilayered complex envelope (Fig. 4A, supplemental movie S2). Consistent with earlier reports, the viral membrane was decorated with HA and NA spikes (Fig. 4B)^14,19,20^. Beneath the lipid bilayer, primary M1 matrix formed a contiguous helical scaffold that remained intact even at the curved filament ends (Fig. 4C). In empty filaments, the characteristic striated pattern of helical M1 was especially clear (Figs. 4D, 3J, 3K). Fourier transform layer-line analysis revealed a helical rise of 3.1 nm corresponding to the repetitive spacing between neighbouring M1 subunits (Fig. 4E), consistent with published values^19^. Notably, some filaments displayed an additional inner helix concentric with M1, sharing the same axis (Figs. 4F, 4G). The majority of these were 1-start helices with inner diameters of 31.6–39.9 nm, outer diameters of 40.1–46.2 nm, helical turn spacing of 7.4-15.9 nm and an average pitch of ∼12 nm (Fig. 3J). In our data, the spacing between strands of the inner helix (12–16 nm) was significantly greater than that of M1 (3.8 nm), corresponding to three or four M1 turns in between, with visible connections to the M1 lattice (Figs. 3J, 3K). A minority of filaments contained 2-start inner helices with a pitch of ∼20 nm and ∼7.6 nm spacing between consecutive strands. Rarely, helices transitioned along the filament length from 2-start to 1-start (only 2 filaments in our data). Inner helices were not always continuous, at times showing interruptions along the filament axis. Similar multi-layered helical assemblies have been reported previously *in vitro* and were postulated to be concentric M1 layers, formed in purified recombinant M1 under high-ionic-strength conditions^20^. Related structures have also been observed in budding virions following disruption of membrane-proximal M1 lattice^53^, where such additional layers could arise from local disengagement of M1 helices. In contrast, in our tomograms of native budding filaments, the membrane-associated M1 lattice appears to remain intact and tightly coupled to both the viral membrane and neighbouring subunits. However, the molecular identity of this additional inner helix remains uncertain, and it may reflect M1 in association with viral or host factors, or an alternative host-derived component.

**Figure 4.**
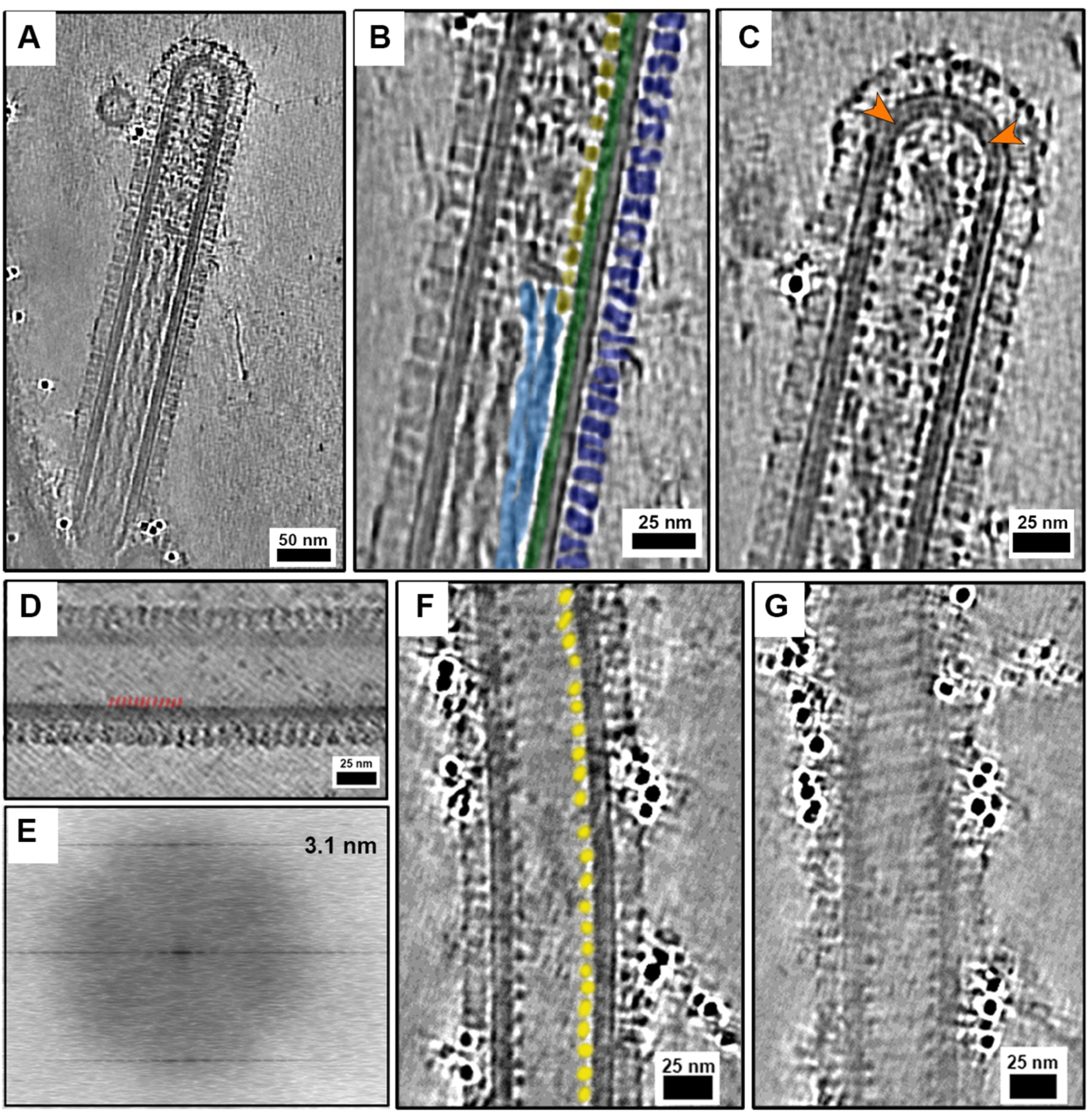
Cryo-ET uncovers complex ultrastructure of IAV filaments. **(A)** High-resolution tomograms showed the complex organisation of filaments composed of glycoproteins, lipid envelope, helical M1 matrix and additional structural features. **(B)** A central slice through a denoised tomogram shows dense packing of glycoproteins (blue). Beneath the lipid bilayer (grey) and helical matrix M1 (green) are highlighted. Additional internal features such as fibrils (light blue) and a second helix (yellow) are also shown. **(C)** M1 is seen to be continuous along the filament length and curved ends (orange arrowheads), with its **(D)** helical pattern clearly visible as striations (red lines) indicating the repeating turns of the helix. Highlighted red lines denoting M1 show only a sub section of the filament for easy visualisation. **(E)** Layer line analysis of tomographic 2D projected images indicated a helical rise of 3.1 nm, corresponding to the minimum distance between neighbouring M1 subunits. **(F)** Many filaments also contained a secondary inner helix sharing the same axis as M1 but with larger spacing (yellow circles) between consecutive helical turns, **(G)** typically arranged as a 1-start helix. Images of the inner helix in F and G are shown along different tomographic Z-slices of the same IAV filament.

Many filamentous virions also enclosed fibrillar densities resembling twisted ribbons with regular constrictions along their length (Fig. 4B). Although we previously reported the presence of this feature in filaments^14^, the improved image quality due to modern cryo-EM detectors has led to substantially improved resolution of fine features in these data. Further, this study focused on straight filament segments rather than filament ends, therefore RNPs were rarely observed (Fig. 3L).

Together, our data demonstrate that native budding IAV filaments are structurally heterogeneous, incorporating distinctive features such as concentric inner helices and internal fibrils. Some filaments contained both unique structures, while others displayed only one structural arrangement (Figs. 3H, 3I, 4B, 4C, 4F, 4G).

### Subtomogram averaging identifies internal fibrils as cofilactin

Having observed internal fibrils within IAV filaments, distinct from the additional helix, we aimed to determine their identity. The number of fibrils varied between filamentous virions: some contained only one or two, while others (33%) harboured three or more (Fig. 5A). Based on morphology, we hypothesised these fibrils were actin. To test this, we analysed layer-line spacings in power spectra of projection images of fibrils extracted from tomograms using SPIDER. These showed a repeating helical pitch of 27.6 nm (supplementary Fig. S2A), differing from the 37 nm pitch of bare actin but correlating well with cofilin-decorated actin, or cofilactin (Figs. 5A, 5B)^55^. To verify this, we applied subtomogram averaging (STA) to resolve 3D structures of fibrils and determine their molecular composition. Volumes containing helical segments were extracted, aligned, and averaged using RELION-5 in STA mode, yielding maps with features consistent with published cofilactin structures^55,56^. Most fibrils corresponded to cofilactin, although close inspection of the tomograms revealed some narrower fibrils resembling bare actin. Executing RELION 3D classification separated fibrils into two distinct classes: cofilactin and actin (Figs. 5A, 5B). After iterative rounds of refinement and averaging, we obtained final maps at 12 Å for cofilactin (Figs 5C, 5D, S2B, supplemental movie S3) and 24 Å for actin (Figs 5E, 5F, S2C, supplemental movie S3). Our data show that cofilin binding shortens the actin helical pitch by ∼27% (from ∼37 nm to ∼27 nm) and alters its geometry (helical rise: 27.9 Å, twist: -166.7°), with cofilactin exhibiting a helical rise of 28.2 Å and twist of –162.3°, consistent with previous reports^55^.

**Figure 5.**
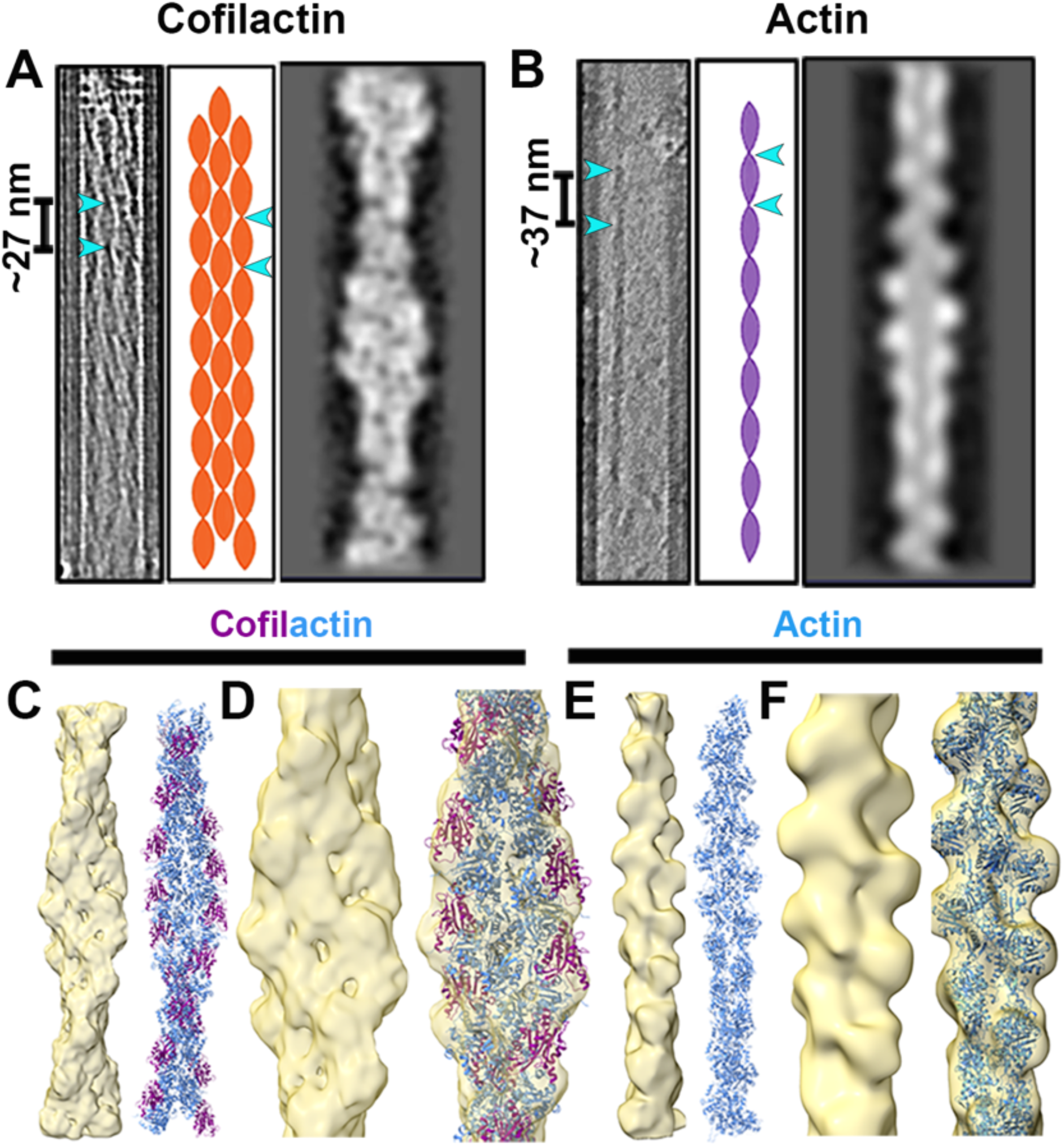
Subtomogram averaging of interior fibrils within IAV filaments suggests they are cofilactin. Structural analysis by STA identified two classes of helical fibrils within IAV filaments: cofilactin and actin. Three panels depicting the tomographic slice, filament illustration and 2D class averages of **(A)** cofilactin and **(B)** actin are shown. Helical pitches of cofilactin (∼27 nm) and actin (∼37 nm) are highlighted. Fitting of AlphaFold-predicted canine **(C)** cofilactin and **(E)** actin into the STA maps demonstrated good fits and confirmed good agreement for both as seen in the **(D, F)** magnified views.

### AlphaFold-aided generation of the canine cofilactin model

Since our STA maps indicated that the fibrils corresponded to cofilactin, we set out to build an atomic model for confirmation. Cofilin, a member of the actin-depolymerising factor (ADF) family, regulates actin dynamics and exists as two isoforms: cofilin 2 in muscle tissue and cofilin 1 in non-muscle cells, such as the MDCK cells used in our study.

Because we infected MDCK cells with IAV, the packaged actin and cofilin observed in our STA structures were of canine origin. BLAST analysis using human actin and cofilin 1 sequences against the dog genome (*Canis lupus familiaris)* revealed 100% identity with their canine counterparts (supplementary Figs S3A,

S3B). AlphaFold3 prediction of the canine actin–cofilin complex closely matched the published rabbit actin–human cofilin filament (PDB 6uc4; supplementary Fig. S3C)^56,57^. Moreover, canine and rabbit actin sequences showed 93% identity (Fig. S3). Using AlphaFold models of canine actin and cofilin, we built extended cofilactin and actin helices in ChimeraX (Figs 5C, 5E) and docked them into our STA maps (Figs 5D, 5F, supplemental movie S3). The strong fit supports our conclusion that the fibrillar density inside IAV filaments is cofilactin.

### Fibrils form higher-order helical bundles of cofilactin within IAV filaments

To investigate how cofilactin fibrils are arranged inside IAV filamentous virions, we mapped the STA maps of cofilactin onto the original sub-tomogram positions within a virion containing 3 fibrils aligned approximately along the z-plane of the tomogram (Figs. 6A, 6B, supplemental movie S3). Notably, cofilactin assembled into tightly packed bundles in approximately 33% of tomograms containing fibrils. Detailed analysis of these bundles revealed that fibrils positioned near the matrix layer and lipid envelope were not strictly parallel but instead displayed curvature in both 2D and 3D views (Figs. 6C, 6E), suggesting a possible long-range right-handed helical organisation. In contrast, fibrils located deeper within the filament, farther from the membrane, remained straight and parallel, lacking visible curvature (Fig. 6D). Overall, our findings indicate that cofilactin fibrils can assemble into higher-order bundles within IAV filaments, potentially stabilising filament structure and coordinating virion assembly, highlighting a functional interplay between cofilactin organisation, actin dynamics and viral budding.

**Figure 6.**
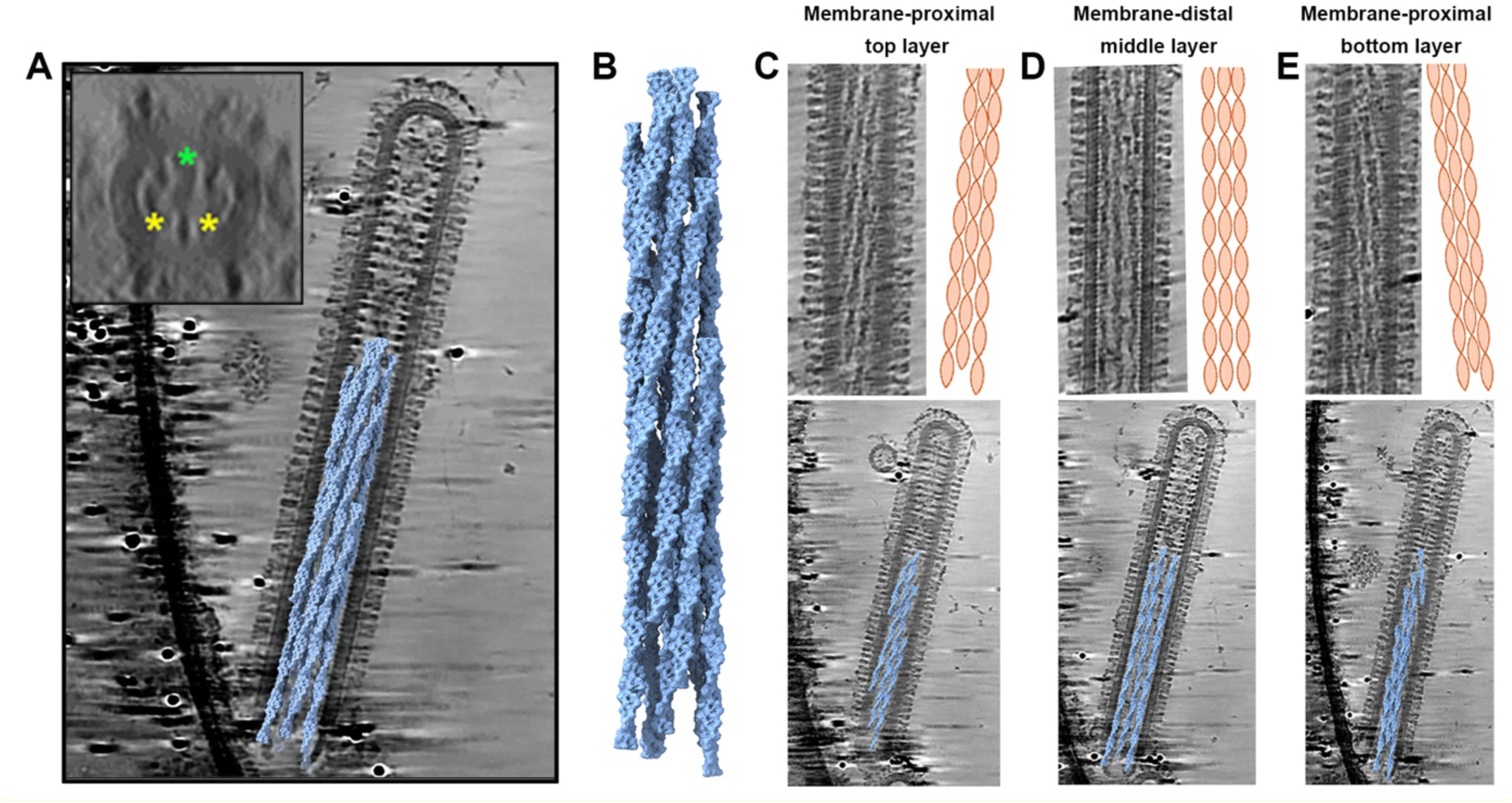
Cofilactin fibrils assemble into helical bundles within IAV filaments. **(A)** Mapping the structure of cofilactin in tomograms indicated **(B)** clustering of cofilactin into tightly packed bundles. Longitudinal sections through the z-plane of the tomogram showed **(C, E)** membrane-proximal cofilactin fibrils near the matrix layer (top and bottom of the virion) were curved in both 2D and 3D, whereas **(D)** membrane-distal fibrils in the middle layer remained straight and parallel. Top panels in C-E show include tomographic slices and schematics illustrating fibril curvature across layers. The inset in (A) shows a transverse section with outer curved (yellow stars) and inner straight (green star) fibrils visible near and away from the membrane, respectively.

### IAV modulates cofilin levels and activity during infection

Given the presence of cofilactin within IAV filaments, we next examined how cofilin expression and localisation are regulated during infection. Confocal microscopy and western blot analyses were performed on A/Udorn/72-infected and mock-treated MDCK cells (Fig. 7). Cofilin is a central regulator of actin turnover, and its ability to bind and sever actin filaments is governed by phosphorylation at serine 3 - phosphorylation inactivates cofilin, while dephosphorylation restores activity^58^. Light microscopy showed IAV filaments budding from actin-rich regions of the plasma membrane, whereas cofilin displayed uniform cytoplasmic distribution, with no marked differences in localisation between infected and mock control cells (Figs 7A). However, quantitative analysis revealed a 1.9-fold increase in total cellular cofilin in infected cells compared with uninfected controls (Fig. 7B). We further confirmed these findings by Western blots of cofilin 1, phosphorylated cofilin-1 (p-cofilin 1) and β-actin normalised to housekeeping protein GAPDH (Fig. 7C). While β-actin levels remained unchanged, infected cells exhibited higher total cofilin 1 expression and a marked reduction in p-cofilin 1 levels (Fig. 7D).

**Figure 7.**
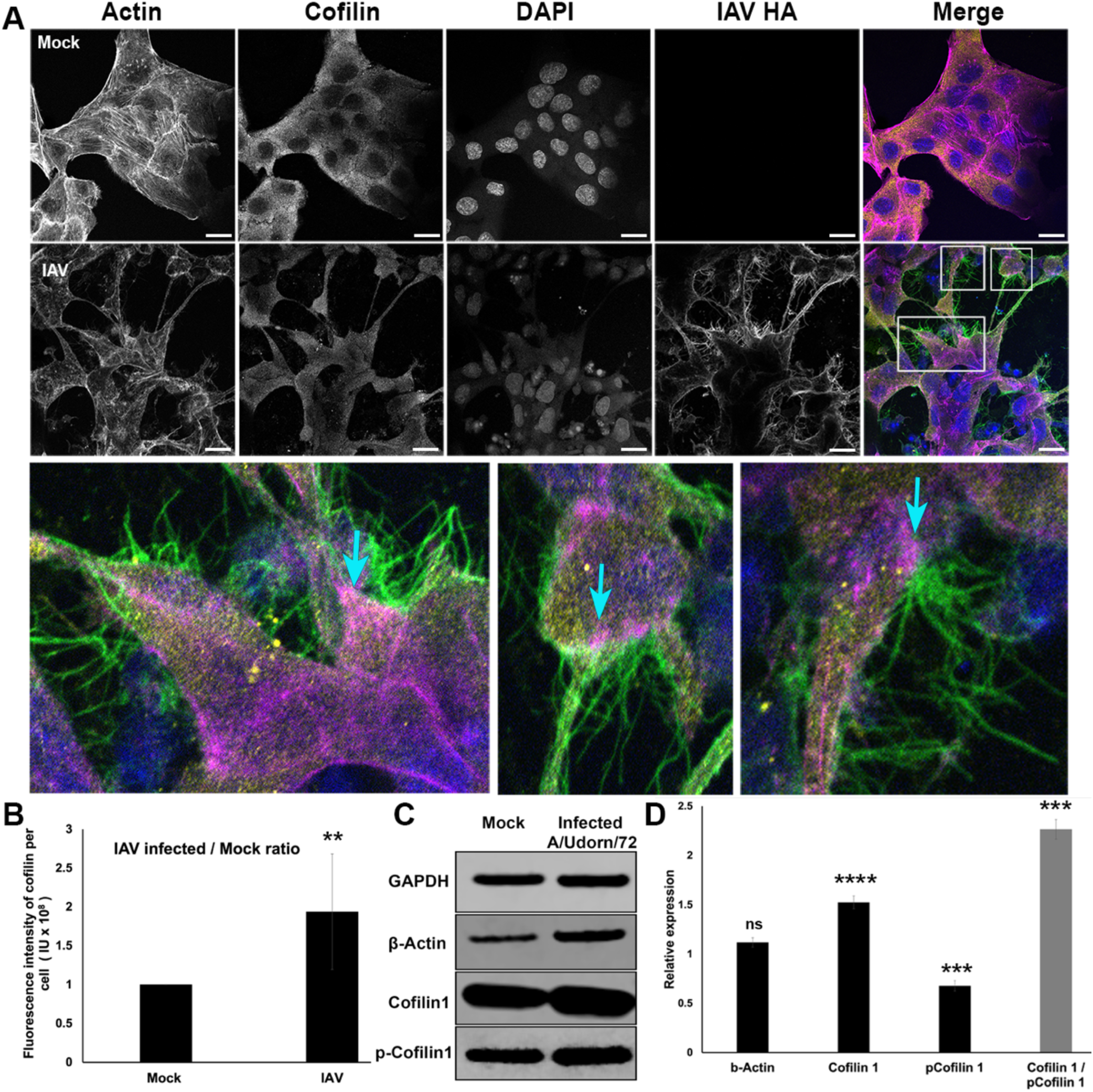
IAV regulates cofilin expression and phosphorylation. MDCK cells were infected with A/Udorn/72 (MOI 0.5) and analysed 24 h post-infection by **(A–B)** confocal microscopy and **(C–E)** western blotting to assess cofilin expression. **(A)** Mock-infected (top) and IAV-infected (bottom) cells were stained for actin (pink), cofilin (yellow), nuclei (DAPI, blue), and viral hemagglutinin (HA, green). Representative single-plane images are shown. IAV filaments bud predominantly from actin-rich regions at the plasma membrane, while cofilin remains evenly distributed throughout the cytoplasm, with no apparent localisation differences between infected and control cells. Scale bar, 20 μm. Enlarged regions (white boxes in A) highlight actin-rich zones of viral egress (cyan arrows). **(B)** Quantitative fluorescence analysis revealed a 1.9-fold increase in total cellular cofilin intensity in infected cells compared to mock controls. **(C)** Western blots show β-actin, cofilin 1, and phosphorylated cofilin 1 (p-Cofilin 1), normalised to GAPDH. **(D)** Relative levels of infected over mock, normalised to GAPDH and mock controls indicated no change in β-actin levels, but a significant increase in total cofilin 1 and a concurrent decrease in pCofilin 1 upon infection. The elevated cofilin 1/pCofilin 1 ratio indicates an accumulation of active, dephosphorylated cofilin during IAV infection. Data represent mean ± SD from three independent experiments (ns P > 0.05; ** P < 0.05; *** P < 0.005; **** P < 0.0001).

The resulting increase in the cofilin 1/p-cofilin 1 ratio indicates an accumulation of active, dephosphorylated cofilin bound to actin in infected cells (Fig. 7D). Collectively, these results demonstrate that IAV infection enhances cofilin expression and activation, consistent with the idea that modulation of cofilin’s actin-binding activity may be a factor in the formation and budding of filamentous virions.

### Integrative model reveals the molecular architecture of IAV filaments

While previous studies have resolved individual components of IAV virions such as RNPs or glycoproteins^11,15-17,21,23^, an integrated high-resolution model capturing the complete molecular organisation of IAV has been lacking. To bridge this gap, we combined our structural and compositional data to construct the first comprehensive integrative model of IAV virions including filaments (Fig. 8; supplemental movie S4). Model construction was guided by the cryo-ET data and reflect the diverse morphologies visualised in our tomograms.

**Fig. 8.**
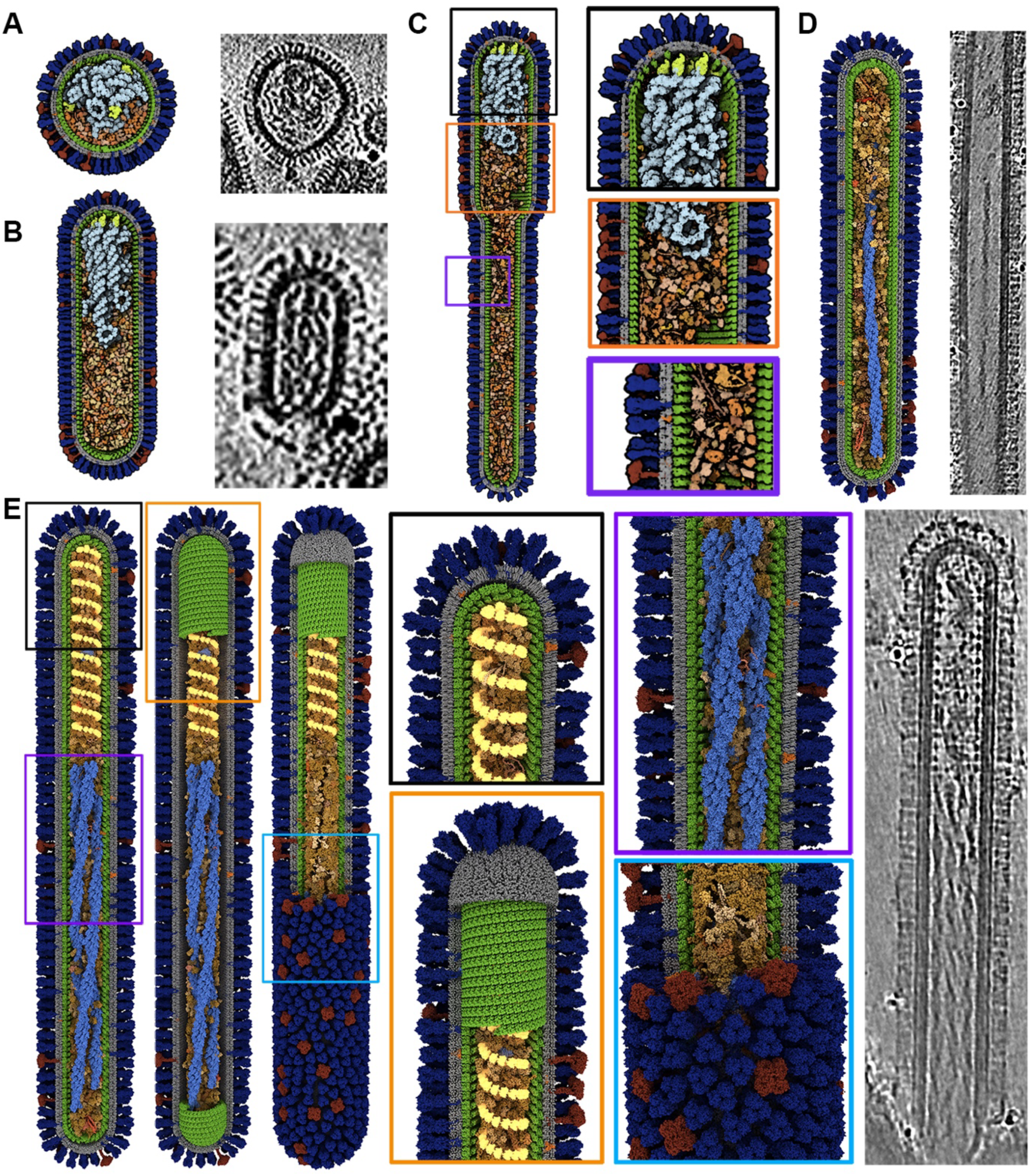
Integrative model of pleomorphic IAV. Comprehensive 3D models depicting the diverse morphologies of influenza A virions and their molecular organization. For reference, comparable morphologies with matching structural features from our tomograms are shown alongside. **(A)** Spherical virions containing unstructured RNPs. **(B)** Bacilliform virions displaying the characteristic “7+1” arrangement of RNP segments. **(C–E)** Filamentous virions illustrating structural diversity, including filaments containing **(C)** RNPs, **(D)** a single cofilactin fibril, or **(E)** bundled cofilactin fibrils. Enlarged views of key features are highlighted by black, orange, and purple boxes in panel (C), and by black, orange, purple, and light blue boxes in panel (E). Models incorporate viral proteins, RNPs, host proteins, cofilactin fibrils and an additional inner helical layer, and are accessible via the **Mol\*** Mesoscale Viewer (https://molstar.org/me/viewer/). Components are colour-coded as follows: HA (dark blue), NA (red), lipid envelope (grey), M1 (green), inner helix (yellow), RNPs (sky blue), polymerase (light green), cofilactin (light blue), and host proteins (shades of beige and brown).

Our integrative modelling approach builds upon previously developed methods^60-62^ where we represent the virion envelope in different geometries upon which the M1 helical lattice was built. Specifically, virion envelopes were modelled using capsule and sphere primitives, both defined as signed distance fields. For each shape, a specific approach was developed to populate the M1 helix along each primitive. In this way, For, each geometry, we developed a tailored procedure to generate the M1 helix on the corresponding surface. This allowed us to cover the different envelope forms with an M1 helix exhibiting the same properties as the helix observed in PDB ID 6Z5J (M1 Y axis, and Z up the membrane ∼3 nm between 2 consecutive M1, and ∼4 nm between two layers). For elongated filament models, we built a second layer of the M1 helix corresponding to the additional inner helix by adding an offset from the primitives and adjusting the number of turns along the main axis to match the distance observed in our tomograms (∼12 nm). Other surface proteins were placed randomly on the envelope.

The 8 vRNPs were modelled using a simplified strategy based on a previously reported approach^63^. The helical region was built from PDB ID 9GAT^64^ (double helix with rise of 24.3 Å and a twist of 57.4°), onto which PDB structure 9C4H^65^ was superimposed as individual NP–RNA units. The loop region was derived from PDB ID 2WFS^66^, again using PDB structure 9C4H, while the polymerase from PDB IS 6RR7^67^ was positioned at the opposite end. The RNA is discontinuous and does not represent a complete full-length sequence. The 8 vRNPs were then arranged in a 7+1 configuration, with seven surrounding a central vRNP, consistent with previously reported organisation^13,68^.

Finally, cofilactin was either generated via a random walk of a single fibril or constructed using a precomputed bundle of helices based on the STA map. The remaining empty space within virions was randomly populated with human host proteins identified from our proteomics data. Structure for each protein were obtained from the Uniprot-AlphaFold database^69^. The complete virion model was then subjected to a simple overlap relaxation using NVIDIA Flex. When the RNPs were not under stress they preserved their shape. Under stress, however, the naive flex implementation failed to maintain correct RNA–protein interactions, although the overall volume was conserved. This is illustrated in supplemental movie S4, where we demonstrate our compression approach for fitting vRNPs inside the small spherical virion. The sphere mesh is first scaled up, then gradually scaled down to bring all vRNP rigid bodies inside the virion. The resulting modelled dataset captures the pleomorphic nature of IAV, encompassing spherical (Fig. 8A), bacilliform (Fig. 8B), and filamentous morphologies (Figs. 8C–E). Notably, filamentous models reveal distinct structural populations—some containing RNPs (Fig. 8C), others cofilactin alone (Fig. 8D), and a subset featuring both cofilactin and the additional inner helix (Fig. 8E). Altogether, these integrative models provide the most complete structural representation of IAV virions and filaments to date and constitute a valuable resource for exploring viral morphogenesis and function. All models are exported as mmCIF files accompanied by manifest files facilitating integration into Mol* Explorer for interactive exploration and community sharing^70^.

## Discussion

We combined cryo-ET with multi-omics approaches to elucidate the structure and composition of the clinically relevant filamentous form of IAV. Lipidomic analysis revealed that filamentous particles are depleted in PE and PI, lipids known to influence membrane curvature and budding. Cryo-ET imaging of intact, budding virions directly on EM grids preserved native architecture and revealed striking structural diversity within filamentous IAV. Alongside the ordered glycoprotein and M1 layers, we observed unique structures such as additional inner helices and fibrillar densities within the filament interior, that were absent in spherical and bacilliform virions. Consistent with earlier reports, Fourier analysis of IAV filaments devoid of internal density confirmed a 3.1 nm spacing between adjacent M1 subunits^14^. Notably, 36% of the filaments displayed multi-layered helical architectures, featuring an additional inner helix, aligned coaxially with the outer M1 layer. Although previous studies suggested such inner helices arise only upon M1 disruption or under high-ionic conditions in recombinant preparations^20,53^, our data reveal this to be a native feature of budding filaments. While M1 has been proposed as the primary driver of filament stability and assembly^19^, our findings raise the possibility that the inner helix may provide an additional structural role in filament architecture. Whether the inner helix consists solely of M1 or M1 in complex with viral or host proteins and understanding its mechanism of assembly during filament formation remains to be determined.

Cryo-ET revealed fibrillar densities within IAV filaments resembling twisted ribbons and using STA we identified these as cofilin-decorated actin (cofilactin). Our 3D reconstructions revealed actin filaments, decorated by cofilin monomers bound to every actin subunit. Consistent with previous findings cofilin binding shortens the helical pitch of actin from 37 to 27 nm^55,56^. The interior fibrils were predominantly cofilactin, with few showing bare actin. Fitting the atomic model of canine cofilactin into our 3D reconstruction confirmed the orientation of cofilin relative to actin within individual fibrils and their ordered arrangement in bundles, where curvature of membrane-proximal cofilactin fibrils suggests long-range helical organisation shaped by the geometry of the underlying matrix layer. Furthermore, Udorn-infected cells showed increased cofilin levels corroborating previous observations^46^, alongside reduced phosphorylated cofilin, indicating that dephosphorylated, active cofilin predominates during IAV infection and remains bound to actin—consistent with our structural observations.

Previous studies have demonstrated that cofilin plays an important role in actin reorganisation during virion assembly and egress of spherical strains of IAV^46^. However, in our cryo-ET data cofilactin fibrils were not detected in spherical and bacilliform virions, and our integrative models suggest that there is unlikely to be sufficient space in the lumen of these virions to contain extended cofilactin fibrils. This suggests that they are a characteristic feature of the filamentous IAV morphology. It is notable that in lipid raft–rich filopodial regions, which have been implicated as sites of IAV filament budding, actin filaments have been observed to bundle and undergo dynamic turnover, with elongation at barbed ends and cofilin-mediated disassembly at pointed ends near the membrane^71^.

Our observations raise the possibility that IAV infection may be associated with localised cofilactin formation at lipid raft–rich filopodial regions, potentially contributing to the structural environment of sites of viral budding. We hypothesise that changes in cofilin activity during IAV infection, regulated by increased dephosphorylation might promote the formation of cofilactin fibrils or bundles beneath the plasma membrane (Fig. 9, supplemental movie S4), but further experimental work would be needed to test this. Future work resolving structures of cortical actin at IAV budding sites on the plasma membrane will be important for understanding these mechanisms and the directionality of cofilin binding to actin.

**Fig. 9.**
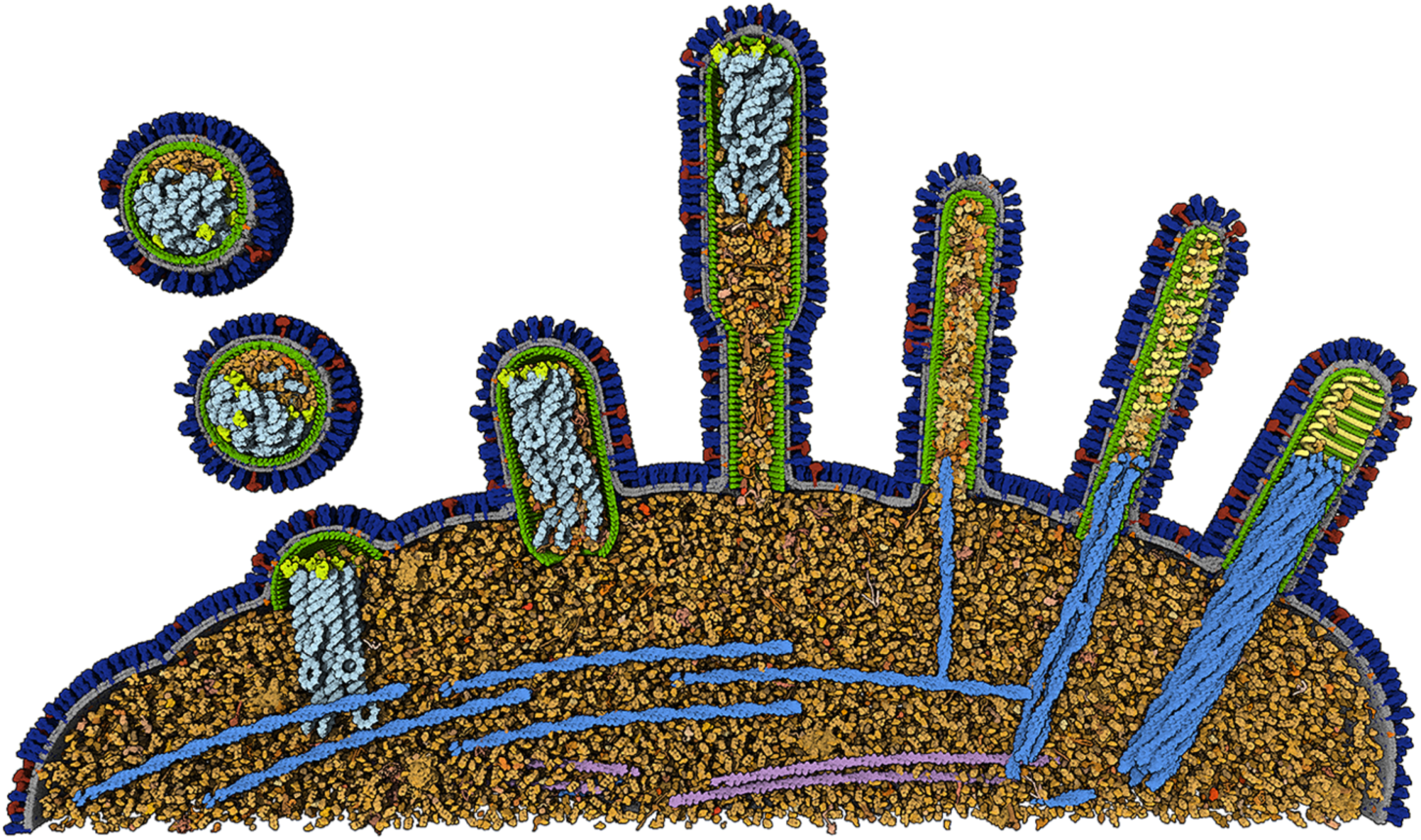
Three-dimensional model of Influenza A virion budding summarizing findings in this study. Illustration of budding IAV virions highlighting distinct architectures of spherical, bacilliform and filamentous particles. Budding occurs at the plasma membrane (grey) where the viral glycoproteins HA (blue) and NA (red) accumulate for recruitment into nascent virions. RNPs (sky blue) are arranged in an ordered configuration within bacilliform particles but appear compact and unstructured in spherical virions. M1 (green), trafficked from the cell interior, assembles at the membrane to form the M1 lattice and drive incorporation of RNPs, ultimately closing at the budding neck prior to virion release. The dense cytoplasmic region beneath the plasma membrane contains several host proteins (shades of beige and brown) including actin filaments (pink) and ribosomes (maroon) that are incorporated into budding virions. Budding filaments are heterogeneous: some package RNPs, whereas others contain internal fibrils and an additional inner helix (yellow). We propose that IAV-induced increases in dephosphorylated cofilin promote localised assembly of cofilactin (light blue) fibrils and bundles beneath the membrane, providing structural support for filament elongation and facilitating budding. The model is accessible via the **Mol\*** Mesoscale Viewer (https://molstar.org/me/viewer/).

By integrating our structural and compositional analyses, we have constructed a detailed integrative model of the molecular architecture of influenza virions and viral filaments. Using cryo-ET and multi-omics approaches, we show that intact IAV filaments pack cofilactin internal fibrils and an additional inner helix and are depleted in lipids important for membrane curvature. Our data reveal that IAV infection induces elevated dephosphorylated cofilin levels and promotes the formation of cofilactin fibrils and bundles, which are subsequently incorporated into budding filaments. This suggests a regulatory role for cofilactin in filament assembly. Collectively, our data enable us to present the first integrative structural model of IAV filaments, laying the foundation for mechanistic studies of their assembly and function.

## Acknowledgements

We thank Prof Andrew Carter (MRC Laboratory for Molecular Biology, Cambridge) for valuable discussions about cofilactin and Dr Chang-Shung Tung (Los Alamos National Laboratory) for advice on the structure of influenza RNPs. We thank Diamond Light Source for access to Cryo-ET facilities at the UK national electron bio-imaging centre (eBIC; proposal EM16637-10), and Dr. Daniel Clare for his support and assistance with tomography data collection at eBIC. We also thank MRC-University of Glasgow Centre for Virus Research Bioimaging for access to light microscopy facilities.

This work was supported by funding from the United Kingdom Medical Research Council to DB and EH [MC_UU_12014/7, MC_UU_00034/1 and MC_UU_00034/7, MR/N008618/1 and MR/V035789/1]; JCH [MC_ST_CVR_2019] and Prof David Robertson [MC_UU_00034/5]. LA was supported by the United States of America National Institute of Health (GM120604 and 5U54AI170855). WK was funded by a European Union’s Horizon 2020 Marie-Sklodowska-Curie fellowship (842067).

## Author Contributions

SV was involved in conceptualisation, formal analysis, investigation, methodology, data curation, visualisation, wrote original draft and reviewed and edited. JCH performed, investigation and methodology, formal analysis and reviewed and edited the manuscript. SC, SSH, CL and TKS performed investigation and reviewed and edited the manuscript. VBS and WK performed formal analysis and reviewed and edited the manuscript. RF was involved in supervision and reviewed and edited. LA was involved in formal analysis, methodology, visualisation, reviewing and editing the manuscript. DB was involved in conceptualisation, funding acquisition, project administration, supervision, visualisation and reviewed and edited the manuscript. EH was involved in conceptualisation and funding acquisition, formal analysis, visualisation, project administration, supervision, writing of the original draft, review and editing.

## Materials and methods

### Cells and Viruses

Madin-Darby canine kidney (MDCK) cells grown in Dulbecco’s modified Eagle’s medium (DMEM) supplemented with 10% foetal calf serum (FCS) were infected with the H3N2 A/Udorn/72 influenza strain at various multiplicities of infection (MOI). For virus purification experiments, an MOI of 0.001 was used, while cryo-ET, light microscopy and western blots utilised an MOI of 0.5. After incubating virus-infected cells at 37°C for 1 h, the media was replaced with serum-free DMEM containing 2.5 µg/ml N-acetyl trypsin (NAT) followed by incubation at 37°C. Virus samples were removed at specific hours post-infection (pi) based on the requirement of the experiment.

### Purification of Influenza virions for multi-omics studies

Influenza virions were purified as described in Hirst and Hutchinson^50^. Briefly, the influenza virus A/Udorn/307/72 (H3N2) was grown in MDCK cells for 96 h. Media were then clarified by low-speed centrifugation and separated by density ultracentrifugation on a 20–30% iodixanol gradient (prepared from OptiPrep in NTC: 0.1 M NaCl, 0.2 M Tris–HCl pH 7.4, 50 mM CaCl_2_) and overnight by ultracentrifugation at 4 °C at 210,000× *g.* Samples from each fraction were adsorbed to electron microscopy grids, negatively stained with 1% phosphotungstic acid and imaged on a JEOL 1200 transmission electron microscope at 20,000× magnification, analysed by Western blot or prepared for omics analyses.

### Purification of Influenza virions for cryo-ET

Following Udorn infection on MDCK cells at MOI 0.001, the supernatant was harvested 36 h pi and clarified by centrifugation at 2500 rpm for 5 min and 10,000 rpm for 30 minutes. Viral supernatant was transferred onto a 30% sucrose cushion in NTE buffer (1 mM EDTA 10 mM, 150 mM NaCl, Tris-HCl, pH 7.5) and pelleted for 2.5 h at 25,000 rpm. Subsequently, the pellet was resuspended in 350 µl NTE and run through a discontinuous sucrose gradient (30%–60%) at 25,000 rpm for 2.5 h. Banded virus was collected and concentrated by centrifuging at 31,000 rpm for 2 h. The pelleted virus was resuspended in 100 µl of NTE buffer.

### Lipidomics

Lipid extractions were performed on fractions 3 and 10 at room temperature, whereby a 1:2 (v/v) ratio of chloroform:methanol (375 µl)was added to the aqueous viral solution (100 µl) within a glass vial, Internal standard, SPLASH® LIPIDOMIX® Mass Spec Standard (Avanti Polar Lipids,) was added to each sample for normalisation. Chloroform and ultrapure water were added to give a final chloroform:methanol:water ratio of 2:2:1 (v/v), and the samples were centrifuged at 1,000 g for 5 minutes at room temperature. The lower phase was transferred to a new glass vial and dried under nitrogen for subsequent lipid analysis.

Lipid extracts were analysed by ESI-MS/MS using a Absceix 4000 QTrap, a triple quadrupole mass spectrometer with a nanoelectrospray source. The dried lipid extracts were reconstituted in 1:2 (v/v) chloroform:methanol and 6:7:2 (v/v) acetonitrile:isopropanol:water, delivered via a nanomate and analysed in positive and negative ion modes using a capillary voltage of 1.25kV. Tandem mass spectra scanning (daughter, precursor, and neutral loss scans) were performed using nitrogen as the collision gas, with collision energies between 35-90 V. Assignment of phospholipid species was based upon a combination of survey, daughter, precursor and neutral loss scans; the phospholipid peaks obtained from the tandem MS/MS were identified and verified using the LIPID MAPS Lipidomics Gateway (http://www.lipidmaps.org). Each phospholipid class was normalised to the corresponding internal standard. The samples were also analysed to obtain high accuracy m/z using a Thermo Exploris Orbitrap mass spectrometer by direct infusion in both positive and negative ion modes.

### Proteomics

Samples were digested for proteomic analysis as previously described^72^. Briefly, 8 M urea, in 1:10 ratio, was added to the samples to a final concentration of 7.2 M urea and denatured for 30 min at room temperature. In final concentrations - 10mM TCEP (Tris(2-carboxyethyl)phosphine) and 50mM iodoacetamide (IAA) were added and incubated in the dark for 30 min at room temperature. Samples were transferred onto FASP filters (filter-aided sample preparation, Vivacon 500, Sartorius, VN01H02, 10kDA) and spun at 14,000 x g, followed by two 200 µl washes with 50 mM TEAB (Triethylammonium bicarbonate). Trypsin (1 µg) in 50mM TEAB (Promega, 1:20 ratio of enzyme : protein) was added and incubated at 37^°^C overnight. The following day, samples were eluted with 200 µl 0.1% trifluoroacetic acid (TFA), followed by 50% acetonitrile (ACN) in 0.1% TFA. Samples were dried in a centrifugal evaporator and re-suspended in 5% DMSO (Dimethyl sulfoxide) in 5% formic acid (FA) prior mass spectrometer analysis.

Samples were analysed by LC-MS/MS using a Vanquish Neo UHPLC (Thermo Fisher) connected to an Orbitrap Ascend Mass Spectrometer (Thermo Fisher). The Vanquish Neo was operated in “Trap and Elute” mode using a PepMap Neo trap 100 (100 μm x 2 cm, Thermo Fisher) and EASY-SPRAY PepMapNeo column (50 cm x 75 μm, 1500 bar, Thermo Fisher). Tryptic peptides were trapped and separated using a 60 min gradient (from 2% to 18% solvent B (0.1% formic acid in acetonitrile) in 40 min and then to 35% B in 20min, flowrate: 300 nl/min).

MS data were acquired in a data-dependent mode (DDA). Briefly, MS1 scans were collected in the orbitrap at a resolving power of 120K, with a scan range (*m/z*) 380-1500. MS1 normalised AGC was set to “Standard” with a maximum injection time set to “Auto” and RF lens at 30%. MS2 scans were then acquired using the ddMS^2^ IT HCD mode with normalised HCD collision energy set at 30% and detector type set to Ion Trap. The Ion Trap scan rate was set to “Rapid” and first mass *(m/z)* at 120. Maximum Injection Time (ms) was set at 50. Ion intensity threshold of 5.0e3 was used. Charge states 2 to 6 were included and the exclusion duration (s) set at 40.

### Mass Spectrometry data analysis

For relative quantification across fractions, iBAQ values were normalized using the R package VSN (v3.50.0)^73^. For each comparison, missing-value imputation was applied only to proteins that were undetected in all replicates of one experimental condition but detected in at least two replicates of the other condition. Imputation was performed using the local minimum determination (“MinDet”) method^74^. Statistical analysis of the processed protein intensities was carried out using the R package limma (v3.38.3)^75^, employing the empirical Bayes–moderated *t*-test, with *p*-values corrected for multiple testing using the Benjamini–Hochberg procedure. The *Homo sapiens* orthologues for *Canis familiaris* proteins were retrieved from InParanoid database^76^.

### Propagation of IAV on cryo-EM grids

MDCK cells seeded on cryo-electron microscopy (cryo-EM) grids were infected with Udorn as previously described^14^. Briefly, glow-discharged 200 mesh gold quantifoil grids (R2/2, Quantifoil Micro Tools GmbH, Germany) were sterilized with ethanol and coated with 50 μg/ml of laminin overnight in glass-bottomed dishes (MATTEK Corporation Inc, USA). Following washes with DMEM, grids were seeded with 10^5^ MDCK cells overnight in DMEM with 10% FCS at 37°C. The following day infection was carried out with Udorn at MOI of 0.5 in serum free DMEM with 2.5 µg/ml NAT and incubated for a further 20 h at 37°C. Grids were then frozen by plunge freezing for cryo-EM.

### Cryo-plunge freezing of grids

Cryo freezing of IAV samples, both purified virus and IAV-infected cells on grids was carried out as previously described^14^. Briefly purified IAV was mixed with 10 nm colloidal gold (BBI Solutions, UK) in a ratio of 1∶3 v/v. An aliquot of 5 µl was applied to glow-discharged Quantifoil 200 mesh copper grids (R2/2, Quantifoil Micro Tools GmbH, Germany) prior to freezing. IAV-infected cells grown on gold Quantifoil grids were supplemented with 5 nm colloidal gold added at a ratio of 1∶3 v/v. All grids were then blotted and frozen by plunging into liquid ethane using a Vitribot Mk IV (Thermo-Fisher Scientific) or Leica EM GP2 (Leica Microsystems).

### Cryo tomogram acquisition and processing

Tomography of purified IAV samples was carried out as described previously^14^. Tilt series of cryo frozen grids of virus-infected cells suitable for subtomogram averaging was collected at the UK Electron Bioimaging Centre (eBIC, Oxford) using a Titan Krios (Thermo-Fisher Scientific) equipped with a Gatan BioQuantum K2 direct electron detector. Dose-symmetric tilt series were recorded using SerialEM^77,78^ at a magnification of 81000, corresponding to a pixel size of 1.773 Å/pixel between −60° to +60° with 3° interval and a defocus between -1 to -3 µm. For each tilt-series, a total of 41 images were recorded with an electron exposure of 2 electrons/Å^2^ per image, and a total of 82 electrons/Å^2^ per tilt-series.

Tomograms were generated and visualized using IMOD^79^. Tomographic reconstruction of purified IAV samples was performed using weighted back projection and figures were prepared by averaging 5 tomogram sections using IMOD’s 3dmod slicer routine. For IAV-infected cellular samples, movies were corrected for beam induced motion using MotionCor2^80^ and then ordered into the final tilt-series stack file using in house perl scripts, SerialEM metadata files (mdoc), and the IMOD command newstack^79^. Tilt-series alignment and reconstruction was carried out by weighted back projection using IMOD^79^ followed by defocus estimation using CTFFIND4^81^. Tomograms were binned by a factor of four and denoised using Topaz to aid visualisation and data interpretation^82^.

### Sub-tomogram averaging

The IAV filament dataset was processed using the subtomogram averaging pipeline of RELION-5^83^. Raw data were imported into RELION-5 and the .mdoc files were renamed as TS[number].mdoc to ensure compatibility for the RELION scripts used later in the processing workflow. Raw data were then imported into RELION-5, and individual tilt movies were motion-corrected and averaged using whole-frame alignment in the RELION implementation of MotionCor2^80,83^. Contrast transfer fraction (CTF) estimation was carried out using CTFFIND-4.1^81^. Nine tilt series were manually inspected and unsuitable tilt images were removed using a Napari plug-in (https://github.com/napari/napari/blob/main/CITATION.cff) provided as part of the ‘exclude tilt images’ job type in RELION-5. Tilt series were then automatically aligned using fiducial-based alignment with IMOD implemented within RELION-5 and tomograms were reconstructed at a pixel size of 10 Å for visual inspection. Denoised tomograms were generated to aid particle picking using the cryo-CARE wrapper in RELION-5 by training 9 tomograms and 1200 sub-volumes per tomogram. Particles were picked from denoised tomograms using the ‘filament picking mode’ within the Napari plug-in as multiple points describing a curve with the Z-axis aligned along the helical axis of the fibrils at an inter-particle spacing of 50 Å. Subtomograms were binned by a factor of 4 and extracted at a pixel size of 7.092 Å/pixel and used to generate an initial 3D reference model with initial values of helical rise and twist of 28.5 Å and -162°, respectively for further processing. Upon reaching maximum Nyquist resolution, subtomograms were then extracted at a binning factor of 2 and refined using the coordinates, orientations and the 3D reference of the bin 4 set of particles. Duplicate particles at a minimum inter-particle distance of 20 Å were then removed prior to classification. Poorly aligned particles were removed by running a 3D classification for five classes and an angular sampling interval of 1.8 degrees using helical parameters of 25-31 Å rise and -160° to -170° twist. Classes corresponding to cofilactin and actin were pooled together and unbinned subvolumes were extracted and a reference map at bin 1 produced using the “Reconstruct particle” job type in RELION-5. Having generated masks in RELION-5, duplicate particles were removed and unbinned subvolumes of cofilactin (1570 particles) and actin (132 particles) were subjected to further auto-refinement using mask and helical parameters. Following 3D refinement, the map was sharpened by postprocessing to estimate the gold-standard FSC curves that generated the 3D reconstruction for actin at 24 Å resolution. For cofilactin, data were further subjected to a single round of particle-specific CTF refinement, followed by extraction of the improved subvolumes, reconstruction of a new 3D reference map and 3D refinement resulting in an EM map at 12 Å resolution. STA maps were visualised using ChimeraX and overlaid on tomograms using ArtiaX extension in ChimeraX^84,85^.

### Sequence alignment

Using sequences of human actin (NCBI ID: NP_001092.1) and human cofilin 1 (NCBI ID: NP_005498.1) as references, protein BLAST search was carried out against the canine genome (*Canis lupus familiaris*) and revealed orthologous sequences of canine actin (NCBI ID: NP_001182774.2) and cofilin 1 (NCBI ID: XP_038280889). Multiple sequence alignment of actin (human, canine and rabbit [NCBI ID: NP_001095153.1]) and cofilin 1 (human and canine) was carried out using Mafft with BLOSUM62 protein scoring matrix and default parameters indicating a sequence identity of 100% between the human and canine sequences (cofilin 1 and actin) and 93% between human and rabbit actin^86^. Sequences were visualised with the R programming language’s MSA package^87^.

### Model building

Canine sequences of actin and cofilin 1 obtained from the BLAST analysis was used to predict the structure of a monomeric canine actin-cofilin unit using Alphafold3^57^. This monomeric cofilactin complex closely resembled both in sequence and structure an earlier PDB structure of cofilactin filament made of rabbit actin and human cofilin (ID: 6uc4) and therefore validated using the Alphafold structure to build our canine cofilactin model. Using a tetramer of two actin-cofilin units, consecutive cofilin-actin units were generated and aligned using the “Matchmaker” tool in ChimeraX, with duplicating subunits being removed^85,88^. Repeating this procedure for 10 consecutive cofilin-actin dimers, single cofilin and actin subunits at either end were removed and the cofilactin filament model assembled. Similarly, the actin filament was built based on the Alphafold structure of canine actin. Assembled cofilactin and actin filament models were fit into the EM reconstructions using ChimeraX^85^.

### Fourier layer line analysis

Layer lines analysis was carried out using SPIDER^89^. Binned tomograms (bin 4 at pixel size of 7.092 Å/px) of empty IAV filaments or filaments containing internal fibrils were trimmed in IMOD and saved as 3D stacks with “mrc” extension. The 3D tomograms were opened in SPIDER and first converted to the spider format with ‘spi’ extension. 2D projections of the 3D stacks were then made using the ‘pj 3’ operation. Projections were padded in a box of 972 pixels along both X and Y dimensions. Calculating the inner and outer half-widths, gaussian soft masks of 9 pixels were applied to the 2D padded projections on all sides along the X and Y directions using the ‘Ma X’ and ‘Ma Y’ operations. Power spectra and their log values were calculated for each filament with the ‘pw’ and ‘pw l’ operations, respectively. Sums of the power spectra and log power spectra were calculated by adding individual power spectra or their log values for 9 IAV filaments with cofilactin and 5 empty IAV filaments using the ‘ad s’ operation. 1D plots of the summed power spectrum along the Y-axis was generated using Bsoft^90^ and imported into Microsoft Excel to plot and measure layer lines.

### Confocal light microscopy

Confluent monolayers of MDCK cells grown on cover slips were infected with Udorn as previously described. Briefly, uninfected control and infected cells at an MOI of 0.5 were fixed with 4% formaldehyde/2.5% Triton X-100 24 h pi. Labelling was then performed by incubating cells with primary antibodies for 1 h at room temperature, followed by washing to remove unbound antibodies and subsequent incubation with fluorescent-tagged secondary antibodies for 30 min at room temperature. Cells were labelled for H3 haemagglutinin (HA) with a mouse monoclonal against A/X-31/1968 H3N2 virus, kindly provided by Prof. John Skehel (NIMR, London), and for cofilin and p-cofilin with rabbit polyclonals (Abcam UK, ab42824 and ab12866) and detected using goat anti-mouse Alexa Fluor 633 (Invitrogen, UK) and donkey anti-rabbit Alexa Fluor 488 (Life Technologies, UK), respectively. Phalloidin Alexa Fluor 568 (Life technologies, UK) was used to stain actin. Cells were then washed and mounted using ProLong Gold Antifade with DAPI (Thermofisher, UK). Normalised 16-bit fluorescent images of cell monolayers were acquired in *.czi format with a Zeiss LSM 880 confocal microscope fitted with a 63x/1.4 Plan-Apochromat oil objective lens (Carl Zeiss). For quantitative analysis, total cofilin intensity across each image was quantitated using ZEN software (Carl Zeiss) and cell numbers were calculated by counting DAPI-stained nuclei. The ratio of cofilin intensity/cell was compared between infected cells and uninfected cells, with > 200 cells counted per population.

### Western blots

Cell lysates of Udorn-infected (MOI 0.5, 24h pi) and uninfected MDCKs were run on NuPAGE 4-12% Bis-Tris gels (Thermo Fisher Scientific) and transferred for 10 minutes onto 0.2 μm nitrocellulose membranes for Western blot analysis using the Trans-blot turbo transfer system (Bio-Rad). Transferred blots were then blocked with 5% milk for one hour at room temperature and subsequently probed with specific antibodies against mouse GAPDH (Thermo Fisher Scientific), mouse β-actin (Cell Signalling), rabbit cofilin 1 (Abcam) and rabbit phosphorylated cofilin 1 (Abcam) overnight at 4°C. After three rounds of washing with PBST (10 mM Na_2_HPO_4_, 1.8 mM KH_2_PO_4_, 137 mM NaCl, 2.7 mM KCl, 0.1% Tween) blots were probed with fluorescent anti-rabbit DyLight 800 and anti-mouse DyLight 680 secondary antibodies (Cell Signalling) for 1 hour at room temperature. Bound antibodies were visualised using the LI-COR Odyssey XF imaging system (LI-COR) and quantified using the Image Studio Lite 5.2 (Li-COR). Experiments were carried out as biological triplicates (n=3 independent experiments), and graphs were plotted and statistically analysed using the two-tailed t-test in Excel. Statistical significance is indicated as ns (P>0.05), *** (P<0.005), **** (P<0.0001).

### Integrative modelling

Virion envelopes were modelled using capsule and sphere primitives defined as signed distance fields. For each shape, a specific approach was developed to populate the M1 helix along each primitive. In this way, different shapes were covered with an M1 helix that exhibits the same properties as the helix observed in PDB ID 6Z5J (M1 Y axis, and Z up the membrane ∼3 nm between 2 consecutive M1, and ∼4 nm between two layers). For elongated models, we built a second layer of the M1 helix corresponding to the additional inner helix by adding an offset from the primitives and adjusting the number of turns along the main axis to match the distance observed in EM (∼12 nm). Other surface proteins are placed randomly on the envelope, while the vRNP positioned inside the virus was built with CA beads, based on an established model^63^, which are then replaced by the actual PDBs for NP (2q06) and polymerase (6RR7) and placed within the different virion shapes. Finally, cofilactin was either generated via a random walk of a single fibril or constructed using a precomputed bundle of helices based on the STA map. The remaining empty space was randomly populated with human host proteins identified from our proteomics data. Structure for each protein was obtained from the uniprot-alphafold database. The complete virion model was then subjected to a simple overlap relaxation using NVIDIA Flex. When RNPs were not under stress they kept their shape, but under stress the naive flex implementation did not maintain proper rna-protein interaction although volume was maintained. All integrative models were exported as mmCIF files and can be easily visualized via Mol* Mesoscale Explorer^70^ and are publicly available to interactively explore (https://molstar.org/me/viewer/). For the summary figure (Fig. 9), because of the large number of proteins, only the surface proteins were relaxed. The number of internal proteins is currently prohibitive for our approach to relax and remove steric clashes.

