## Supplementary Figures S1 to S3 for "Molecular architecture of Influenza A virions"

Supplementary data

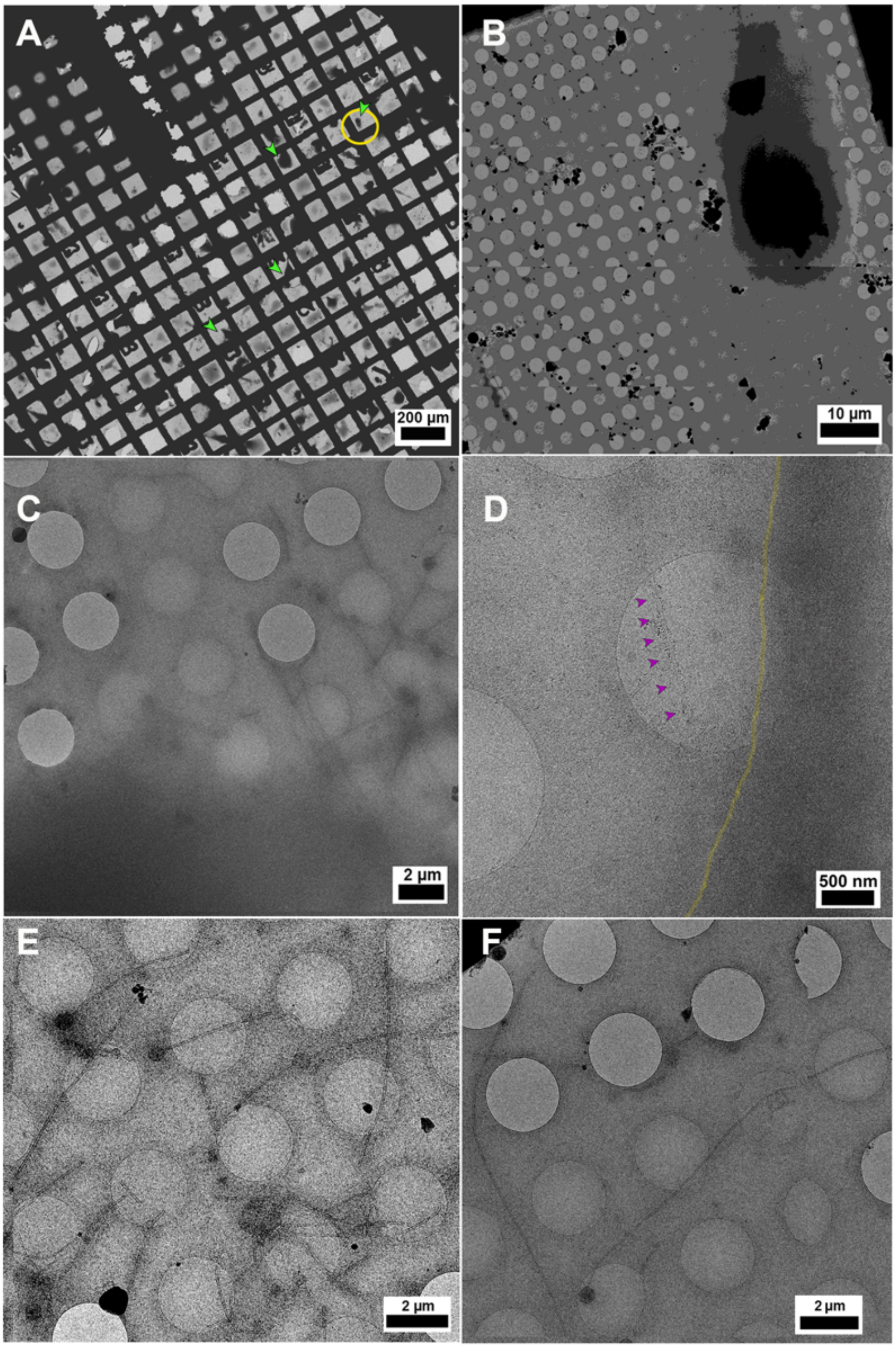


**Supplementary Figure S1. Cryo-electron tomography of budding filamentous IAV particles budding from plunge frozen cells.**

Cells were propagated and infected on gold Quantifoil (R2/2) EM grids and imaged for IAV filamentous virion budding by cryo-ET. **(A)** Montages of complete grids at low magnifications showed good ice thickness and cell dispersity (green arrowheads) (A). **(B)** Magnified image of the cell in (A) annotated as yellow circle is shown as an example where regions of interest for tilt series collection were targeted on areas that had good vitrification around its edges. Imaging infected cell edges at low magnifications **(C)** revealed several intact filaments in the vicinity including **(D)** filamentous virions (pink arrowheads) in the process of budding from the cell edge (yellow line). **(E)** Budding filaments were well preserved and often **(F)** extremely long (> 5-12 mm).

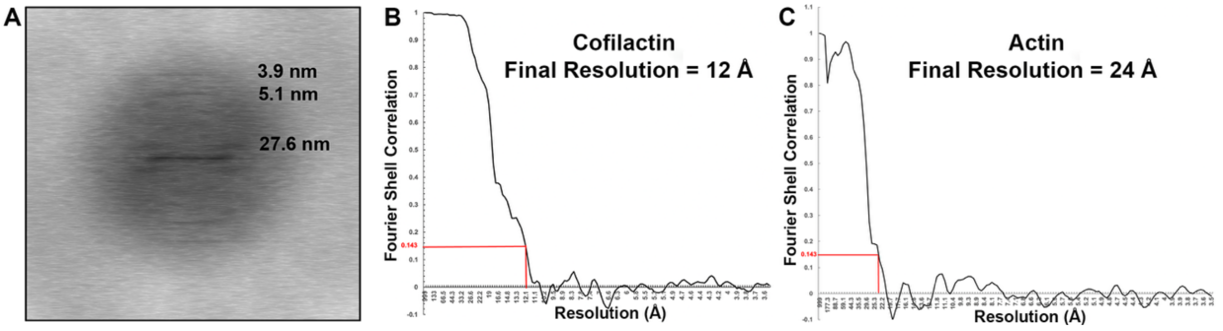


**Supplementary Figure S2. Helical parameters and resolution of internal fibrils inside IAV filamentous particles.**

**(A)** Layer line analysis showed that fibrillar density within filaments had a helical pitch of 27.6 nm corresponding to the distance between repeating turns of the helix. FSC curves from STA 3D reconstructions of fibrils resolved structures of **(B)** cofilactin at 12 Å and **(C)** actin at 24 Å at FSC 0.143.

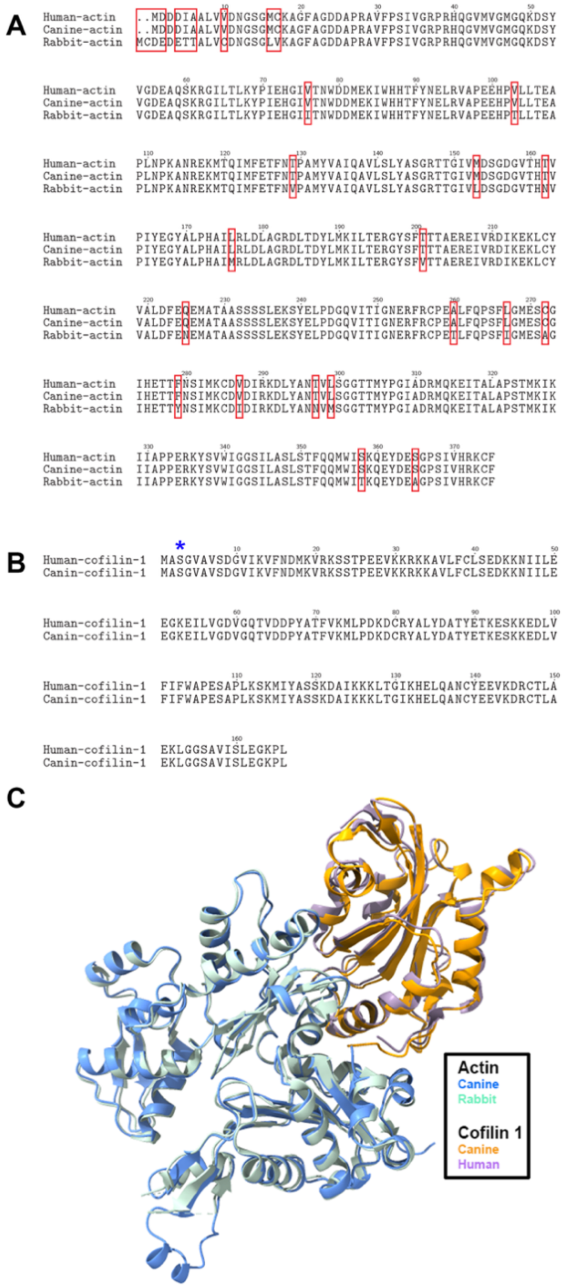

**Supplementary Figure S3. Sequence and structural conservation of actin and cofilin.**
**(A)** Protein BLAST sequence analysis showed 100% identity between human and canine actin, while rabbit and canine actin shared 93% identity with residue differences highlighted (red boxes). **(B)** Human and canine cofilin 1 were also identical, including the phosphorylation site at Ser3 (blue star). **(C)** Structural comparison of rabbit actin–human cofilin (PDB 6uc4) with the AlphaFold-predicted monomeric canine actin–cofilin complex revealed strong similarity, validating use of the AlphaFold model as a template for building our cofilactin atomic model.
